# Efficacy of a new multivalent vaccine for the control of bovine respiratory disease (BRD) in commercial fattening units

**DOI:** 10.1101/2024.08.07.606972

**Authors:** M. Tapiolas, M. Gibert, C. Montbrau, E. Taberner, M. Solé, H. Santo Tomás, A. Puig, R. March

## Abstract

Bovine respiratory disease (BRD) is the most common cause of morbidity and mortality in cattle. The effects of BRD are most marked during the first weeks after arrival in calf-rearing units. Vaccination is a tool that can help to control the disease by reducing both the incidence itself and the massive use of antibiotics. Recently, a new multivalent vaccine (DIVENCE®), containing live gE/tk double-gene deleted BoHV-1, live-attenuated BRSV, inactivated PI3, BVDV-1 recombinant protein and BVDV-2 recombinant protein, has been designed to protect cattle against the main viral pathogens associated with BRD. The aim of this study was to demonstrate the efficacy of DIVENCE® against BRD in field conditions. A total of 360 animals from three different batches were included in the study: one batch of 108 Holstein-Friesian males (Farm 1), another batch of 99 Holstein-Friesian males (Farm 2), and 153 Belgian-Blue cross males and females (Farm 3). On each feedlot, a single batch of animals was included in the study, with a mean age of approx. ten weeks (73.4±0.6 days). On the vaccination day (Day 0 of the study; D0), calves from the same batch were randomly distributed between the two study groups. The vaccinated group (n=183) received the DIVENCE® vaccine and the control group (n=177) received a placebo injection of phosphate-buffered saline (PBS) solution. Both groups were given two intramuscular doses (2 mL/dose) of the corresponding product three weeks apart. All animals were monitored during the entire fattening period (approx. 9 months) after vaccination to assess the incidence, severity, and morbidity of BRD as well as administered treatments and feed performance. Overall, vaccinated animals had significantly (*p*<0.004) lower morbidity than controls, with only 49 out of the 183 vaccinated calves (26.78%) presenting at least one episode of respiratory disease (RD), versus 73 out of the 177 control calves (41.24%). Thus, a reduction of 35.1% in morbidity was observed in the vaccinated animals. In terms of RD cases, vaccinated animals had a significantly (*p*<0.001) lower number of cases than control animals, with only 66 cases reported in the 183 vaccinated calves (0.36 cases/calf) versus 110 cases in the 177 control calves (0.62 cases/calf). A reduction of 42% in cases was observed in the vaccinated animals. During the study follow-up, a BRSV outbreak was reported in one of the farms (Farm 1) on Day 23, just two days after the second dose. The vaccinated group had significantly (*p*<0.02) lower morbidity (11 animals out of 54; 20.4%) and severity (score of 1.70) compared to the control group (29 animals out of 54; 53.70% and score of 2.11). Overall, vaccinated animals needed significantly (*p*=0.01) fewer antimicrobial treatments (at least one due to RD) than controls. Furthermore, vaccinated animals presented numerically higher average daily weight gain (*p*=0.07) and significantly higher carcass weight (*p*=0.01) than controls, at 35.78 g/day and 6.58 kg, respectively.

Vaccination with DIVENCE® at the beginning of the fattening period decreased the incidence and morbidity of BRD following a BRSV outbreak only two days after the primary vaccination scheme. Additionally, the incidence and morbidity of BRD throughout the entire fattening period was reduced in the vaccinated animals. Thus, DIVENCE® can improve economic outcomes in fattening units by reducing antibiotic treatments and enhancing performance.

## INTRODUCTION

Bovine Respiratory Disease (BRD) is the most common cause of morbidity and mortality in cattle (Holman et al., 2015). This disease is a multifactorial syndrome mostly associated with bovine respiratory syncytial virus (BRSV), infectious bovine rhinotracheitis (IBR), bovine viral diarrhea virus (BVDV), and bovine parainfluenza virus 3 (PI-3). All these viruses are known to be involved alone or in combination with each other and/or with bacteria in the pathogenesis of BRD (Fulton, 2009); BRD is often the result of interactions between stress and bacterial and viral infections. In feedlots, BRD is usually observed in the first period of the fattening phase when animals must cope with stressful events such as transportation, commingling, and weaning (Timsit et al., 2011; Schneider et al., 2009).

The negative economic impact of BRD due to the increase in mortality and treatment costs (metaphylactic and therapeutic use of antimicrobials) alongside the reduction in performance and carcass value has been reported by several authors worldwide (Thompson et al., 2006; Cusack et al., 2007; Fulton, 2009; Pardon et al., 2013; Fernández et al., 2020). Furthermore, a high incidence of BRD increases the use of antibiotics, which spurs an increase in antibiotic resistance (Catry et al., 2016). These effects, coupled with the growing concern for animal welfare, create a need to find solutions to improve animal health through other strategies, such as prevention.

The effects of BRD are more marked during the first weeks after arrival in calf-rearing units, when mortality and morbidity rates are usually high (Pardon et al., 2012; Winder et al., 2016). Controlling the BRD complex involves improving both environmental and management conditions while preventing viral and bacterial infections through vaccination (Pratelli et al., 2021, Fulton et al., 2015, Cusack, 2023). One common management practice is to vaccinate young calves early in life to provide immune protection before stressful situations, although vaccination is usually done upon arrival on the feedlot (Jourquin et al., 2023). The objective of these measures is to improve the feedlot health program in order to minimize the incidence and costs associated with morbidity and mortality (BRD and other diseases) through designated prevention and control programs, thus maximizing feeding performance and carcass value (Edwards, 2010). Recently, a new multivalent vaccine (DIVENCE®), containing live gE/tk double-gene deleted BoHV-1, live-attenuated BRSV, inactivated PI3, BVDV-1 recombinant protein and BVDV-2 recombinant protein, has been designed to protect cattle against the main viral pathogens associated with BRD. Efficacy of the vaccine against these viruses has been demonstrated in experimental conditions, with protection against BRD being achieved. The vaccine is also effective for fetal protection in animals challenged with ncp BVDV-1 and BVDV-2 (Taberner et al., 2024). The aim of this randomized field trial was to evaluate the efficacy of DIVENCE® vaccine in young calves on reducing the incidence, morbidity, and need for treatment for BRD while enhancing average daily weight gain (ADWG) on three different commercial fattening units.

## MATERIALS AND METHODS

### Study animals (animals and housing)

The study was conducted on three commercial cattle feedlots located in the northeast region of Spain. On each feedlot, a single batch of animals was included in the study. A batch was considered as the group of calves that arrived at the farm on the same day and remained together for the entire fattening period. All animals enrolled in the study were approx. ten weeks of age (73.4±0.6 days of age), with the minimum age at vaccination being 55 days at Day 0 of the study. No animals had been previously vaccinated against BRSV, BoHV-1 (IBR), PI3, BVDV-1 or BVDV-2. A total of 360 animals from three different batches were included in the study: one batch of 108 Holstein-Friesian males (Farm 1), another batch of 99 Holstein-Friesian males (Farm 2), and 153 Belgian-Blue cross males and females (Farm 3).

Each batch remained housed in the same airspace for the entire fattening period in outdoor pens, open at the front and partially closed at the back. All animals were commingled together throughout the study. All animals were fed *ad libitum* using concentrate and forage to meet nutritional requirements.

### Study design

The study was a controlled, randomized, double-blind, parallel-field trial. On the vaccination day (Day 0 of the study; D0), calves from the same batch were randomly distributed between the two study groups. The vaccinated group (n=183) received the DIVENCE® vaccine and the control group (n=177) received a placebo injection of phosphate-buffered saline (PBS) solution. Both groups were given two intramuscular doses (2 mL/dose) of the corresponding product three weeks apart. The personnel involved in the vaccination procedure conducted no other study-related tasks.

### Clinical observations

Animals were checked daily by the farmer and weekly by the veterinarian from vaccination (D0) until the end of the fattening period, approx. 9 months after vaccination. During this period, animals showing respiratory clinical signs were considered as a ‘case of respiratory disease (RD)’. A score was given to all RD cases using a 4-point scale to evaluate the severity of clinical signs based on previous work from Torres et al., (2013). This score was as follows: 0: clinically normal; 1: mild signs of depression with mild respiratory signs; 2: moderate signs of depression with moderate respiratory distress; 3: severe signs of depression with severe respiratory distress; 4: moribund.

An outbreak was considered if more than 10% of animals showed signs of RD for 2-3 consecutive days following the recommendations described by the Spanish Ministry of Agriculture, Fisheries and Food (MAPA, 2023). Animals were then monitored daily for five days after the outbreak. Nasal exudate samples from the reported cases were also collected to detect specific pathogens by means of PCR.

Antimicrobial metaphylactic treatments were not allowed during the entire study period so that RD could be properly explored. If a calf presented respiratory signs, individual antimicrobial treatment was administered per criteria from the farmer/veterinarian. Individual treatments with respiratory purposes and cases of mortality were recorded during the overall follow-up period. Necropsies were performed by the veterinarian whenever possible.

### Performance evaluation

Days on feed were calculated from D0 to slaughter. Individual live body weight was recorded at D0 and individual carcass weight was obtained from the slaughterhouse. Average daily weight gain (ADWG, g/day) was calculated as the difference between live body weight at D0 and the estimated final body weight in relation to days on feed. The estimated final live body weight was determined considering a 55% dressing percentage (MAPA, 2022).

### Data analysis

Animals that did not complete the vaccination schedule (2 doses, 21 days apart) were excluded from the statistical analysis. The statistical analysis was done using R software v3.1. A *p-*value < 0.05 was considered as the limit for statistical significance. To compare the two groups, four analyses were performed per variable: one for each feedlot and one overall. In the overall analysis, the farm was used as random effect. The overall incidence of RD was analyzed using a Poisson regression model and a Cox’s proportional hazards survival model. Morbidity was analyzed using a logistic regression model, as well as odds ratio and a Cox’s survival model. The number of treatments (one or more antimicrobial treatments) and mortality were analyzed using a logistic regression model. A Cox’s survival model was also performed for mortality. Productivity data (days on feed, weight at Day 0, growth and carcass weight) were analyzed using a t-test on each farm or a linear regression model (overall test).

## RESULTS

### Incidence, mortality, and antimicrobial treatments due to respiratory disease

A total of 360 calves were followed up until the end of fattening. At the beginning of the study (D0), no statistical differences were observed between groups in terms of age and body weight as detailed in Table 1.

**Table 1.** Descriptive statistics of study animals at vaccination day (D0).

|  | Farm 1 |  | Farm 2 |  |
| --- | --- | --- | --- | --- |
|  | Control | Vacc. | Control | Vacc. |
| Animals (N) | 54 | 54 | 48 | 51 |
| Mean age (days) ± SE | 75.2 ± 1.18 | 76.4 ± 1.20 | 76.0 ± 1.67 | 74.4 ± 1.39 |
| *p*-value | 0.34 |  | 0.67 |  |
| Mean weight (kg) ± SE | 92.0 ± 1.76 | 91.1 ± 1.77 | 97.1 ± 1.47 | 95.3 ± 1.35 |
| *p*-value | 0.74 |  | 0.37 |  |
|  | Farm 3 |  | Overall |  |
|  | Control | Vacc. | Control | Vacc. |
| Animals (N) | 75 | 78 | 177 | 183 |
| Mean age (days) ± SE | 69.3 ± 1.29 | 71.7 ± 1.78 | 72.9 ± 0.82 | 73.8 ± 0.93 |
| *p*-value | 0.76 |  | 0.68 |  |
| Mean weight (kg) ± SE | 100.5 ± 2.21 | 105.2 ± 2.32 | 97.0 ± 1.18 | 98.3 ± 1.26 |
| *p*-value | 0.14 |  | 0.34 |  |

During the study period, the percentage of vaccinated animals on Farms 1 and 2 that presented at least one episode of RD was significantly (*p*<0.05) lower compared to morbidity in the control groups (Table 2). Low morbidity was observed in both groups on Farm 3. Overall, vaccinated animals had significantly (*p*<0.004) lower morbidity than controls, with only 49 out of the 183 vaccinated calves (26.78%) presenting at least one episode of RD, versus 73 out of the 177 control calves (41.24%). Thus, a reduction of 35.1% in morbidity was observed in vaccinated animals.

**Table 2.** Prevalence of RD for each individual feedlot and overall data during the follow-up period.

|  | Farm 1 |  | Farm 2 |  | Farm 3 |  | Overall |  |
| --- | --- | --- | --- | --- | --- | --- | --- | --- |
|  | Control | Vacc. | Control | Vacc. | Control | Vacc. | Control | Vacc. |
| At least one case of RD |  |  |  |  |  |  |  |  |
| Number of cases | 31 | 18 | 23 | 14 | 19 | 17 | 73 | 49 |
| % | 57.4 | 33.3 | 47.9 | 27.4 | 25.3 | 21.8 | 41.2 | 26.8 |
| p-value | 0.01 |  | 0.04 |  | 0.61 |  | 0.004 |  |
| Odds ratio | 0.37 (0.17-0.81) |  | 0.45 (0.18-0.95) |  | 0.82 (0.39-1.74) |  | 0.52 (0.33-0.81) |  |
| Total cases of RD |  |  |  |  |  |  |  |  |
| Number of cases | 58 | 24 | 27 | 17 | 25 | 25 | 110 | 66 |
| Cases/calf | 1.07 | 0.44 | 0.56 | 0.33 | 0.33 | 0.32 | 0.62 | 0.36 |
| p-value | <0.001 |  | 0.09 |  | 0.89 |  | <0.001 |  |

Regarding the total number of RD cases reported during the entire study, vaccinated animals on Farm 1 were observed to have significantly (*p*<0.05) fewer cases than controls. On Farms 2 and 3, the number of cases in vaccinated groups was numerically lower than in control groups. Overall, vaccinated animals had significantly (*p*<0.001) fewer cases than control animals, with only 66 cases reported in the 183 vaccinated calves (0.36 cases/calf) versus 110 cases in the 177 control calves (0.62 cases/calf). A reduction of 42% in cases was observed in the vaccinated animals.

During the study follow-up, one respiratory outbreak was reported in one of the farms (Farm 1) on Day 23, just two days after the second dose. Laboratory results confirmed BRSV as the only etiological pathogen responsible for the outbreak. On this farm, during the outbreak follow-up, the vaccinated group had significantly (*p*<0.001) fewer animals affected by RD (11 out of 54; 20.4%) than the control group (29 out of 54; 53.70%). Consequently, a reduction in RD cases of 62.1% in the vaccinated group was observed. The odds of having RD were 0.22 (95% CI: 0.22 [0.09-0.52]) in vaccinated animals versus control animals; control animals are thus 4.5 times more likely to suffer from RD. The severity score of clinical signs during the outbreak was also reported to be significantly (*p*<0.02) lower in the vaccinated group (1.70) than the control group (2.11). During the outbreak, five calves died due to RD. Findings at necropsy showed acute pulmonary emphysema with extensive pulmonary lesions, consistent with BRSV. The mortality rate reported in the vaccinated group was numerically lower than in the control group (1.85% versus 7.41%).

In terms of the antimicrobial treatments reported for RD during the study follow-up, the percentage of animals that required at least one treatment in the vaccinated group on Farm 1 was significantly (*p*=0.01) lower than the control group (33.3% vs 57.4%). On Farm 2, the percentage of calves that required at least one treatment was also lower in the vaccinated group in comparison to control group (27.4% vs 43.7%). On Farm 3, the groups were found to be similar in this regard (22.7% vs 21.8%). Overall, vaccinated animals needed significantly (*p*=0.01) fewer antimicrobial treatments (at least one due to RD) than controls. Similar results were observed for the total number of calves that required more than one antimicrobial treatment for RD (29.6% vs 13.0%, 12.5% vs 3.9%, and 5.3% vs 5.13% for Farms 1, 2 and 3 respectively). Overall, the percentage of vaccinated animals that received more than one antimicrobial treatment (14.7% vs 7.1%) was significantly (*p*<0.02) lower than in control group.

### Productivity parameters

Animals spent an average of 269 days in the study from vaccination until slaughter. The groups were considered homogeneous in terms of initial body weight (*p*=0.34). Performance results are summarized in Table 3. ADWG and carcass weight were found to be greater in vaccinated animals compared to controls on all three farms. Overall, vaccinated animals presented numerically higher ADWG (*p*=0.07) and significantly higher carcass weight (*p*=0.01) than controls, at 35.78 g/day and 6.58 kg, respectively.

**Table 3.** Performance results for each individual feedlot and overall data during the follow-up period (mean ± SE).

|  | **Farm 1** |  | **Farm 2** |  | **Farm 3** |  | **Overall** |  |
| --- | --- | --- | --- | --- | --- | --- | --- | --- |
|  | Control | Vacc. | Control | Vacc. | Control | Vacc. | Control | Vacc. |
| **Days on feed (d)** | 303.7 ± 1.16 | 301.9 ± 1.86 | 265.5 ± 2.45 | 266.5 ± 3.36 | 247.0 ± 2.32 | 248.4 ± 2.17 | 268.5 ± 2.21 | 269.1 ± 2.19 |
| *p*-value | 0.41 |  | 0.82 |  | 0.67 |  | 0.85 |  |
| **ADWG (g/d)** |  |  |  |  |  |  |  |  |
| **Overall** | 1313.5 ± 20.02 | 1345.6 ± 17.17 | 1364.7 ± 15.26 | 1391.3 ± 19.25 | 1357.5 ± 22.97 | 1402.4 ± 22.71 | 1346.8 ± 12.32 | 1382.6 ± 12.28 |
| *p*-value | 0.23 |  | 0.28 |  | 0.17 |  | 0.07 |  |
| **RD** | 1286 ± 30.29 | 1346 ± 32.41 | 1343 ± 20.73 | 1344 ± 40.75 | 1268 ± 45.91 | 1329 ± 41.57 | 1300 ± 18.75 | 1340 ± 21.56 |
| *p*-value | 0.18 |  | 0.98 |  | 0.33 |  | 0.17 |  |
| **No RD** | 1345 ± 24.46 | 1345 ± 20.44 | 1386 ± 22.18 | 1408 ± 21.38 | 1397 ± 25.07 | 1418 ± 26.07 | 1383 ± 15.71 | 1396 ± 14.56 |
| *p*-value | 0.99 |  | 0.47 |  | 0.55 |  | 0.46 |  |
| **Carcass weight (kg)** |  |  |  |  |  |  |  |  |
| **Overall** | 269.6 ± 3.59 | 273.7 ± 3.07 | 252.2 ± 2.26 | 256.0 ± 3.26 | 238.2 ± 2.86 | 247.8 ± 2.47 | 251.1 ± 1.99 | 257.7 ± 1.84 |
| *p*-value | 0.39 |  | 0.33 |  | 0.012 |  | 0.01 |  |
| **RD** | 263 ± 4.81 | 275 ± 5.54 | 249 ± 3.26 | 248 ± 4.49 | 225 ± 5.88 | 236 ± 5.77 | 248 ± 3.26 | 254 ± 3.95 |
| *p*-value | 0.13 |  | 0.78 |  | 0.17 |  | 0.08 |  |
| **No RD** | 277 ± 5.09 | 273 ± 3.74 | 255 ± 2.71 | 259 ± 4.04 | 243 ± 2.97 | 250 ± 2.61 | 253 ± 2.43 | 259 ± 2.06 |
| *p*-value | 0.58 |  | 0.42 |  | 0.079 |  | 0.19 |  |

When ADWG and carcass weight were compared between animals with RD and those without, higher ADWG was still observed in vaccinated animals despite RD. Comparable results were observed for carcass weight, with overall higher results (*p*=0.08) for carcass weight in vaccinated animals with RD than corresponding controls.

## DISCUSSION

BRD continues to be a major problem in feedlots, not only because of the disease itself, but also because of the impairment of productive performance and the high use of antimicrobials. Historically, this high use of antimicrobials as a metaphylactic measure has long been a common practice on fattening farms to control BRD morbidity and mortality. However, the European regulation on the restriction of veterinary drugs (EU 2019/6), which mainly focuses on the use of antibiotics, and WHO guidelines (2017) on use of antimicrobials in food-producing animals, have sought to promote new strategies to prevent and improve disease control, which include vaccination.

The objective of the present study was to evaluate the efficacy of a pentavalent viral vaccine (DIVENCE®) under field conditions, focusing on the assessment of incidence, morbidity and antimicrobial treatments related to BRD and productivity parameters in different feedlots. During the study, one BRD outbreak was observed on Farm 1. This outbreak occurred 23 days after the start of the vaccination program, two days after the second dose (98.7 ± 8.67 days of age). This rapid onset of a BRD outbreak during the first days of fattening is a common occurrence (Edwards 2010, Leruste et al., 2012); in a previous study (Macartney et al., 2003) involving over 12,000 fattening calves, it was found that 80% of cases of BRD were observed within the first 55 days after entering the feedlot. Despite the outbreak occurring immediately after completion of the vaccination program, a significant (*p*<0.01) reduction in incidence was observed in vaccinated animals compared to controls. Laboratory results identified BRSV as the etiological pathogen; lung lesions were also compatible with this. BRSV can be considered as one of the main pathogens in BRD (Makoschey and Berge, 2021; Studer et al., 2021), especially during the first year of life (Marzo et al., 2021, Santo Tomás et al., 2023a). The results obtained demonstrate that the attenuated vaccine virus strain of BRSV contained in DIVENCE® induces a protective immunological response. This vaccine strain has been proven efficacious against an experimental BRSV infection when used intranasally in young animals (Marzo et al., 2021) several weeks after vaccination. The results observed in the present study, when efficacy was clearly observed two days after the second intramuscular dose, suggest that the vaccine induces both a humoral and cellular response, resulting in a rapid response of vaccinated animals to the natural infection. This cellular response was previously described using a vaccine containing the same attenuated BRSV strain (Marzo et al., 2021). The most relevant clinical signs observed during the outbreak were dyspnea followed by nasal discharge and cough; few animals died. The significant differences in clinical signs observed were between vaccinated and control animals, suggesting that the vaccine reduced the severity of the infection.

It is worth mentioning that, although some animals were individually protected and immunized, herd immunity was not achieved in these scenarios, as only 50% of the animals were vaccinated against the main BRD viruses. Considering that for BRSV, BoHV-1 and BVDV, the basic reproductive numbers (R0) in naïve populations are 36.5, 3.2, and 3.4, respectively (de Jong et al., 1996; Bosch et al., 1998, Moerman et al., 1993), herd immunity is not feasible by vaccinating only half of the animals. Some authors (Milián-Suazo et al., 2022) suggest that at least 70% of a herd must be immunized to obtain herd immunity in similar situations. The results are better in controlling viral circulation and the impact of the disease when herd immunity is achieved (Roeder et al., 2007). This suggests that, if at least 70% of the animals had been vaccinated, the outbreak could have been reduced, since it would have minimized the number of affected animals, reducing the total excretion of the herd, and consequently reducing the basic reproductive number. However, despite the circumstances, a decrease in incidence, severity, and even mortality was observed in the vaccinated group during the outbreak. Veal calf mortality has been estimated at approximately 2.9-5.1% in Europe (Studer et al., 2021). The mortality rate in the vaccinated group (1.85%) was lower than that reported in field conditions, while the mortality rate of control animals was higher (7.4%). During the study, the vast majority of RD cases were reported during first months of the follow-up period. Specifically, 52% of RD cases were reported within the first month after initiation of the study, and 43% between first and third month after day D0. Only 6% of RD cases were observed afterwards. These results confirm that most BRD cases occur during the first days of the fattening period, as other authors have described (Edwards 2010, Leruste et al., 2012; Macartney et al., 2003). These findings reveal the importance of prevention and control of respiratory disease during the first months of fattening. Farm 1 suffered the highest incidence and morbidity of RD cases during the study period, because of the respiratory outbreak. Nevertheless, the incidence of RD on Farm 2, where no outbreak was reported, was significantly lower in the vaccinated group (27.4%) compared to the control group (47.9%). Farm 3 exhibited a similar incidence and morbidity across both groups. However, considering all data from all three feedlots, the overall incidence and morbidity of RD cases was significantly lower in vaccinated animals, which suggests that the effect of viruses associated with BRD had a lesser impact on those animals.

This decrease in incidence, severity, and morbidity may favor a reduction in antibiotic treatment. It is well known that mixed infections are frequently identified in BRD and bacteria are BRD-associated pathogens (Santo Tomás et al., 2023b). It also is likely that prior viral infections (BRSV, PI3, IBR or BVDV) can predispose cattle to severe bacterial bronchopneumonia while going undiagnosed due to the overwhelming bacterial lesions (Divers 2018). For this reason, antimicrobials are commonly used once an animal or group of calves present clinical signs of respiratory disease. Antibiotics, however, target bacteria and mycoplasma, and are not effective against viruses. Some authors (Neethirajan et al., 2018) have described BRSV infections actually occurring after an animal is treated with antibiotics for a bacterial infection, which raises concerns about massive antibiotic use and consequent antibiotic resistance. On the other hand, if BRD is not properly diagnosed, its delayed detection can impair the efficacy of the treatment and the expected recovery (Hanzlicek et al., 2010). In many cases, antimicrobial treatment begins when clinical signs are already very evident, so it is very likely that lung damage has already occurred. This can be detrimental to short and long-term health status, because lung consolidation is linked to increased mortality, delayed growth, lower reproductive performance, and shorter life expectancy (Kamel et al., 2024). Prevention is key to improving animal welfare, complying with new legislation that calls for a reduction in the use of antibiotics (i.e., Regulation (EU) 2019/6 in the European Union), and above all minimizing the costs associated with BRD. In the present study, a clear and significant reduction in antibiotic treatment in the overall analysis was observed. This reduction could be explained by vaccine-induced protection against the most relevant BRD viruses. Viral agents, such as BRSV, BHV-1, BVDV, parainfluenza-3 virus, and others, further weaken the immune defenses of calves, paving the way for bacterial infections that lead to serious respiratory issues (Gaudino et al., 2022; Zhang et al., 2019; Fulton et al., 2020). Consequently, the use of the vaccine reduced the incidence and morbidity of the most relevant BRD viruses, preventing bacteria from easily colonizing the respiratory tissue and thereby reducing the number of antibiotic treatments.

The differences observed between vaccinated and control animals had an impact on performance. Overall, higher ADWG (g/d) and carcass weight were observed in vaccinated animals compared to control animals. This result was observed when considering all the animals, only those animals with RD, and only those animals without RD. The negative impact of BRD on performance has already been widely reported. The consequences of BRD in young calves include reduced ADWG (Cuevas-Gómez et al., 2021; Cramer et al., 2019). In animals with BRD during their first month of fattening, ADWG can fall by 370 g/day (Schneider et al., 2009). While there may be compensatory growth afterwards, their total growth compared to the healthy counterparts in the group will not recover. Thus, ADWG in animals with BRD is reduced by 70-90 g/day overall compared to healthy calves (Schneider et al., 2009; Smith 1998). This reduction in ADWG impacts days on feed and/or carcass weight. In addition to the cost of treating animals with BRD, there is another cost that can be overlooked: the extra days spent in finishing by animals that have had BRD (Fernández et al., 2020). This can amount to an additional 5.5 days until slaughter for every four and a half months on feed (Thompson et al., 2006). Apart from the cost of feed or labor, delaying the slaughter of these animals entails an opportunity cost, as new calves could be occupying their places during this time, which means fewer calves finished per year. In the present study, animals in both groups (vaccinated and control) had similar body weight at the beginning of the fattening period and had practically the same number of days on feed. However, the overall average carcass weight was significantly higher in the vaccinated group compared to control animals. This indicates that the performance of vaccinated animals was greater, suggesting that the reduction in BRD induced by the vaccine minimizes the impact on the carcass. It has been widely reported (Pardon et al., 2013; Studer et al., 2021; Fernández et al., 2020) that the carcasses of calves with lung lesions at slaughter weigh less, have worse traits (carcass quality), and arrive after more time on the farm, which translates into long-term cost. It should also be noted that subclinical BRD, without mortality or even associated veterinary costs, can still induce a substantial reduction in ADWG and carcass quality (Griffin, 2014; William, 2007). It is estimated that, at best, only 50% of animals with BRD show respiratory symptoms (Buczinski et al., 2018). On Farm 3, lower incidence and morbidity values were observed compared to the other farms, but vaccinated animals had significantly higher carcass weights compared to controls, suggesting that undetected or subclinical BRD cases impair performance by impacting carcass weight in the long term. Consequently, the use of the vaccine reduced the incidence and morbidity of the most relevant BRD viruses, preventing bacteria from easily colonizing the respiratory tissue, and thereby reducing the number of antibiotic treatments. In summary, by minimizing the impact of BRD on the herd through vaccination, i.e., decreasing the incidence and morbidity of BRD as well as antibiotic treatments, higher performance in terms of carcass weight was achieved.

## CONCLUSION

Vaccination with DIVENCE® at the beginning of the fattening period decreased the incidence and morbidity of BRD following a BRSV outbreak only two days after the primary vaccination scheme. Additionally, the incidence and morbidity of BRD was reduced in vaccinated animals throughout the fattening period. Thus, DIVENCE® can improve the economic outcomes in fattening units by reducing antibiotic treatments and enhancing performance.

## Acknowledgments

The authors would like to thank Marc Vila for the help performing the study as well as all personnel at HIPRA and HIPRA Scientific involved in the study.

## References

- Bosch JC, De Jong MC, Franken P, Frankena K, Hage JJ, Kaashoek MJ, Maris-Veldhuis MA, Noordhuizen JP, Van der Poel WH, Verhoeff J, Weerdmeester K, Zimmer GM, Van Oirschot JT. An inactivated gE-negative marker vaccine and an experimental gD-subunit vaccine reduce the incidence of bovine herpesvirus 1 infections in the field. Vaccine. 1998; 16(2−3):265−271. doi: 10.1016/s0264-410x(97)00166-7.

- Buczinski S, Fecteau G, Dubuc J, Francoz D. Validation of a clinical scoring system for bovine respiratory disease complex diagnosis in preweaned dairy calves using a Bayesian framework. Prev Vet Med. 2018; 156:102−112. doi:10.1016/j.prevetmed.2018.05.004.

- Catry B, Dewulf J, Maes D, Pardon B, Callens B, Vanrobaeys M, Opsomer G, de Kruif A, Haesebrouck F. Effect of Antimicrobial Consumption and Production Type on Antibacterial Resistance in the Bovine Respiratory and Digestive Tract. PLoS One. 2016; 11(1):e0146488. doi:10.1371/journal.pone.0146488.

- Cramer MC, Ollivett TL. Growth of preweaned, group-housed dairy calves diagnosed with respiratory disease using clinical respiratory scoring and thoracic ultrasound-A cohort study. J Dairy Sci. 2019;102(5):4322–4331. doi:10.3168/jds.2018-15420.

- Cuevas-Gómez I, McGee M, Sánchez JM, O’Riordan E, Byrne N, McDaneld T, Earley B. Association between clinical respiratory signs, lung lesions detected by thoracic ultrasonography and growth performance in pre-weaned dairy calves. Ir Vet J. 2021; 74(1):7. doi: 10.1186/s13620-021-00187-1.

- Cusack PM, McMeniman NP, Lean IJ. Feedlot entry characteristics and climate: their relationship with cattle growth rate, bovine respiratory disease and mortality. Aust Vet J. 2007; 85(8):311–316. doi:10.1111/j.1751-0813.2007.00184.x

- Cusack PM. Evaluation of practices used to reduce the incidence of bovine respiratory disease in Australian feedlots (to November 2021). Aust Vet J. 2023; 101(6):230−247. doi: 10.1111/avj.13239.

- De Jong MC, van der Poel WH, Kramps JA, Brand A, van Oirschot JT. Quantitative investigation of population persistence and recurrent outbreaks of bovine respiratory syncytial virus on dairy farms. Am J Vet Res. 1996; 57(5):628−633.

- Edwards TA. Control methods for bovine respiratory disease for feedlot cattle. Vet Clin North Am Food Anim Pract. 2010; 26(2):273−284. doi: 10.1016/j.cvfa.2010.03.005.

- Fernández M, Ferreras MDC, Giráldez FJ, Benavides J, Pérez V. Production Significance of Bovine Respiratory Disease Lesions in Slaughtered Beef Cattle. Animals (Basel). 2020; 10(10):1770. doi: 10.3390/ani10101770.

- Fulton RW, d’Offay JM, Eberle R, Moeller RB, Campen HV, O’Toole D, Chase C, Miller MM, Sprowls R, Nydam DV. Bovine herpesvirus-1: evaluation of genetic diversity of subtypes derived from field strains of varied clinical syndromes and their relationship to vaccine strains. Vaccine. 2015; 33(4):549−558. doi: 10.1016/j.vaccine.2014.11.033.

- Fulton RW. Bovine respiratory disease research (1983-2009). Anim Health Res Rev. 2009; 10(2):131−139. doi: 10.1017/S146625230999017X.

- Fulton RW. Viruses in Bovine Respiratory Disease in North America: Knowledge Advances Using Genomic Testing. Vet Clin North Am Food Anim Pract. 2020; 36(2):321–332. doi:10.1016/j.cvfa.2020.02.004.

- Gaudino M, Nagamine B, Ducatez MF, Meyer G. Understanding the mechanisms of viral and bacterial coinfections in bovine respiratory disease: a comprehensive literature review of experimental evidence. Vet Res. 2022; 53(1):70. doi:10.1186/s13567-022-01086-1.

- Griffin D. The monster we don’t see: subclinical BRD in beef cattle. Anim Health Res Rev. 2014; 15(2):138–141. doi:10.1017/S1466252314000255.

- Hanzlicek GA, White BJ, Mosier D, Renter DG, Anderson DE. Serial Evaluation of Physiologic, Pathological, and Behavioral Changes Related to Disease Progression of Experimentally Induced Mannheimia Haemolytica Pneumonia in Postweaned Calves. Am. J. Vet. Res. 2010; 71(3):359–369. doi:10.2460/ajvr.71.3.359.

- Holman DB, McAllister TA, Topp E, Wright AD, Alexander TW. The nasopharyngeal microbiota of feedlot cattle that develop bovine respiratory disease. Vet Microbiol. 2015; 180(1−2):90−95. doi: 10.1016/j.vetmic.2015.07.031.

- Jourquin S, Lowie T, Debruyne F, Chantillon L, Clinquart J, Pas ML, Boone R, Hoflack G, Vertenten G, Sustronck B, Pardon B. Effect of on-arrival bovine respiratory disease vaccination on ultrasound-confirmed pneumonia and production parameters in male dairy calves: A randomized clinical trial. J Dairy Sci. 2023; 106(12):9260−9275. doi: 10.3168/jds.2023-23438.

- Kamel MS, Davidson JL, Verma MS. Strategies for Bovine Respiratory Disease (BRD) Diagnosis and Prognosis: A Comprehensive Overview. Animals (Basel). 2024; 14(4):627. doi: 10.3390/ani14040627.

- Leruste H, Brscic M, Heutinck LF, Visser EK, Wolthuis-Fillerup M, Bokkers EA, Stockhofe-Zurwieden N, Cozzi G, Gottardo F, Lensink BJ, van Reenen CG. The relationship between clinical signs of respiratory system disorders and lung lesions at slaughter in veal calves. Prev Vet Med. 2012; 105(1−2):93−100. doi: 10.1016/j.prevetmed.2012.01.015.

- Macartney JE, Bateman KG, Ribble CS. Health performance of feeder calves sold at conventional auctions versus special auctions of vaccinated or conditioned calves in Ontario. J Am Vet Med Assoc. 2003; 223(5):677−683. doi: 10.2460/javma.2003.223.677.

- Makoschey B, Berge AC. Review on bovine respiratory syncytial virus and bovine parainfluenza - usual suspects in bovine respiratory disease - a narrative review. BMC Vet Res. 2021; 17(1):261. doi: 10.1186/s12917-021-02935-5.

- Marzo E, Montbrau C, Moreno MC, Roca M, Sitjà M, March R, Gow S, Lacoste S, Ellis J. NASYM, a live intranasal vaccine, protects young calves from bovine respiratory syncytial virus in the presence of maternal antibodies. Vet Rec. 2021; 188(11):e83. doi: 10.1002/vetr.83.

- Milián-Suazo F, González-Ruiz S, Contreras-Magallanes YG, Sosa-Gallegos SL, Bárcenas-Reyes I, Cantó-Alarcón GJ, Rodríguez-Hernández E. Vaccination Strategies in a Potential Use of the Vaccine against Bovine Tuberculosis in Infected Herds. Animals (Basel). 2022; 12(23):3377. doi: 10.3390/ani12233377.

- Ministerio de Agricultura, Pesca y Alimentación. Protocolo de metafilaxia para el complejo respiratorio bovino en vacuno de carne [Internet]. 2023. MAPA. https://www.mapa.gob.es/es/ganaderia/temas/sanidad-animal-higiene-ganadera/documentometafilaxiavacunocarne_tcm30-661702.pdf

- Ministerio de Agricultura, Pesca y Alimentación. Informe de base de datos técnico-económica. Ejercicio económico 2022. [Internet]. MAPA. https://www.mapa.gob.es/es/ganaderia/temas/produccion-y-mercados-ganaderos/informebbdd_vacunoleche_2023_publicacion_web_tcm30-639443.pdf.

- Moerman A, Straver PJ, de Jong MC, Quak J, Baanvinger T, van Oirschot JT. A long term epidemiological study of bovine viral diarrhoea infections in a large herd of dairy cattle. Vet Rec. 1993; 132(25):622−626. doi: 10.1136/vr.132.25.622.

- Neethirajan S, Ragavan V, Weng X. Agro-defense: Biosensors for food from healthy crops and animals. Trends in Food Science & Technology. 2018; 73;25−44.10.1016/j.tifs.2017.12.005.

- Pardon B, De Bleecker K, Hostens M, Callens J, Dewulf J, Deprez P. Longitudinal study on morbidity and mortality in white veal calves in Belgium. BMC Vet Res. 2012; 8:26. doi:10.1186/1746-6148-8-26.

- Pardon B, Hostens M, Duchateau L, Dewulf J, De Bleecker K, Deprez P. Impact of respiratory disease, diarrhea, otitis and arthritis on mortality and carcass traits in white veal calves. BMC Vet Res. 2013; 9:79. doi:10.1186/1746-6148-9-79.

- Divers TJ. Respiratory Diseases. Rebhun’s Diseases of Dairy Cattle. 2008; 79–129. doi:10.1016/B978-141603137-6.50007-7.

- Pratelli A, Cirone F, Capozza P, Trotta A, Corrente M, Balestrieri A, Buonavoglia C. Bovine respiratory disease in beef calves supported long transport stress: An epidemiological study and strategies for control and prevention. Res Vet Sci. 2021; 135:450–455. doi: 10.1016/j.rvsc.2020.11.002.

- Regulation (EU) 2019/6 of the European Parliament and of the Council of 11 December 2018 on veterinary medicinal products and repealing Directive 2001/82/ECOJ L 4: 43−167.

- Roeder PL, Taylor WP. Mass vaccination and herd immunity: cattle and buffalo. Rev Sci Tech. 2007; 26(1):253−263.

- Santo Tomás H, Barreto M, Vazquez B, Villoria P, Teixeira R, Sole M. Bovine Respiratory Disease Complex: Prevalence of the Main Respiratory Viruses Involved in Pneumonia in Spain. J Anim Sci Res. 2023a; 7(1): dx.doi.org/10.16966/2576-6457.163.

- Santo Tomás H, Teixeira R, Chacón G, Lázaro S, Sánchez-Matamorors A, Villoria P. Bovine Respiratory Disease Complex: Prevalence of the Different Bacteria Involved in Pneumonia in the Iberian Peninsula. J Anim Sci Res. 2023b; 7(1): dx.doi.org/10.16966/2576-6457.162.

- Schneider MJ, Tait RG Jr, Busby WD, Reecy JM. An evaluation of bovine respiratory disease complex in feedlot cattle: Impact on performance and carcass traits using treatment records and lung lesion scores. J Anim Sci. 2009; 87(5):1821−1827. doi: 10.2527/jas.2008-1283.

- Smith RA. Impact of disease on feedlot performance: a review. J Anim Sci. 1998; 76(1):272−274. doi: 10.2527/1998.761272x.

- Studer E, Schönecker L, Meylan M, Stucki D, Dijkman R, Holwerda M, Glaus A, Becker J. Prevalence of BRD-Related Viral Pathogens in the Upper Respiratory Tract of Swiss Veal Calves. Animals (Basel). 2021; 11(7):1940. doi: 10.3390/ani11071940.

- Taberner E, Gibert M, Montbrau C, Muñoz I, Mallorquí J, Santo Tomás H, Prenafeta A, March R. Fetal protection against bovine viral diarrhea virus types 1 and 2 after vaccination of the dam with the DIVENCE vaccine. BioRxiv. Preprint posted online April 15, 2024. doi: 10.1101/2024.04.12.589196

- Thompson PN, Stone A, Schultheiss WA. Use of treatment records and lung lesion scoring to estimate the effect of respiratory disease on growth during early and late finishing periods in south African feedlot cattle. J Anim Sci. 2006; 84:488–498. doi:10.2527/2006.842488x.

- Timsit E, Bareille N, Seegers H, Lehebel A, Assié S. Visually undetected fever episodes in newly received beef bulls at a fattening operation: occurrence, duration, and impact on performance. J Anim Sci. 2011; 89(12):4272–4280. doi:10.2527/jas.2011-3892.

- Torres S, Thomson DU, Bello NM, Nosky BJ, Reinhardt CD. Field study of the comparative efficacy of gamithromycin and tulathromycin for the treatment of undifferentiated bovine respiratory disease complex in beef feedlot calves. Am J Vet Res. 2013; 74(6):847–853. doi: 10.2460/ajvr.74.6.847.

- WHO guidelines on use of medically important antimicrobials in food-producing animals. Guideline. 7 November 2017. [Internet]. WHO https://www.who.int/publications/i/item/9789241550130.

- William P, Green L. Associations between Lung Lesions and Grade and Estimated Daily Live Weight Gain in Bull Beef at Slaughter. Cattle Pract. 2007; 15(3):244–249.

- Winder CB, Kelton DF, Duffield TF. Mortality risk factors for calves entering a multi-location white veal farm in Ontario, Canada. J Dairy Sci. 2016; 99(12): 10174– 10181. doi:10.3168/jds.2016-11345.

- Zhang M, Hill JE, Fernando C, Alexander TW, Timsit E, van der Meer F, Huang, Y. Respiratory Viruses Identified in Western Canadian Beef Cattle by Metagenomic Sequencing and Their Association with Bovine Respiratory Disease. Transbound Emerg Dis. 2019; 66(3):1379–1386. doi:10.1111/tbed.13172.

